# Correlation of calcium multiphoton with ultrastructural STED imaging of the slit diaphragm in the same glomerulus

**DOI:** 10.1101/2023.09.27.559730

**Authors:** Eva Wiesner, Julia Binz-Lotter, Agnes Hackl, David Unnersjö-Jess, Nelli Rutkowski, Thomas Benzing, Matthias J. Hackl

**Author notes:** Corresponding author: Matthias J. Hackl. contributed equally (Eva Wiesner, Julia Binz-Lotter).

## Abstract

In recent years functional multiphoton imaging of viable mouse tissue and STED imaging of optically cleared tissues allowed new insights into kidney biology. Here, we present a novel workflow where multiphoton imaging of calcium signals in podocytes is, in the same glomerulus, correlated with super-resolved STED imaging and analysis of the slit diaphragm (SD) morphology. Mice expressing the calcium indicator GCaMP3 exclusively in podocytes served as healthy controls and were challenged with two different doses of nephrotoxic serum (NTS). NTS-induced SD architecture damage increased in a dose-dependent manner, whereas intracellular calcium levels exhibited a wide range of variation and showed no correlation with SD damage in the same glomerulus. Our co-imaging protocol is applicable to a broad range of research questions as it is suitable for other tissues and compatible with a variety of reporters and tissue antigens.

**Translational statement:** The combination of multiphoton and super-resolution microscopy identifies the modulation of signaling pathways and resolves the changes in ultrastructure in disease states in the same glomeruli. Using our new method, we can compare functional and morphological data to investigate the degree of correlation to reveal novel underlying pathomechanism. This knowledge can improve our ability to draw conclusions from human kidney biopsies, in which the ultrastructure will be assessed more-often by routine super-resolution microscopy in the future.

## Introduction

Tight control of intracellular calcium levels, actin cytoskeleton and SD structure in podocytes is crucial for kidney function. Human mutations in the ion channel TRPC6, the actin binding protein α-actinin4 and the SD protein podocin all lead to focal segmental glomerulosclerosis (FSGS), loss of kidney function and end stage kidney disease.^1-3^ Acute increases of intracellular calcium levels in podocytes in cell culture result in a reorganization of the actin cytoskeleton, a process called calcium-mediated actin reset^4^. This calcium-actin-slit diaphragm axis connects calcium signaling to SD morphology and is essential for maintaining podocyte function. *In vivo* multiphoton imaging of calcium levels in podocytes using mice expressing the calcium indicator GCaMP3^5-8^ revealed low basal levels and increased calcium signaling after podocyte injury. In parallel, novel super-resolution, methods have allowed the visualization and quantification of SD damage during different diseases.^9-11^ So far, these functional and structural parameters were obtained in separate experiments, impeding correlative studies in single glomeruli. To technically overcome this shortcoming, we established a novel protocol using acute kidney slices (AKS) for multiphoton calcium imaging followed by STED microscopy in the same slice of tissue, which can be used to address new questions regarding podocyte biology.^12^

## Methods

### Mice

Pod:cre-GCaMP3 (C57Bl6/N) mice of both sexes, age 4 weeks were used. Control mice (n=3) received no treatment; NTS model: single i.p. injection of nephrotoxic serum (NTS, Probetex, PTX-001-S, San Antonio, TX, USA): low dose 8 μl/g (n=4) or high dose 11 μl/g (n=3). No method of randomisation or blinding was used. Mice were bred and kept in IVC under SPF conditions in an in vivo research facility (CECAD, Cologne). Mice were monitored for weight loss and proteinuria starting day 3 (not shown). After 5 days kidneys were harvested after cervical dislocation. All experiments were performed following guidelines provided by the LANUV NRW, Germany approval number: VSG 81-02.04.2018.A351. Animal experiments were reported using ARRIVE guidelines.^13^

### Preparation of acute kidney slices (AKS)

Fresh mouse kidneys were embedded in low-melting agarose and cut into 300 μm slices using a vibratome (LEICA VT1200 S, Mannheim, Germany). Slices were maintained in carbogenated Krebs-Henseleit-Buffer.^14^ AKS were incubated with 5 μM propidium iodide for 5 min to label cells damaged by cutting.

### Preparation of acute kidney slices for STED microscopy

Immunofluorescent labeling was conducted as previously described^9^. Primary antibodies used: guinea pig anti-nephrin (Fitzgerald, 20R-NP002, 1:100), GFP-Alexa647 (Thermo Fisher, A-31852, 1:100). Secondary antibody: donkey anti-guinea pig Atto594 (coupled in house; 1:100).

### Multiphoton and STED microscopy

Multiphoton imaging was performed with a TCS SP8 MP-OPO microscope (Leica Microsystems, Mannheim) with a Chameleon Vision II laser (Coherent) tuned to 940nm. Burning marks for reorientation were created by focusing increased laser power (50% power, 8 s) on small points of tissue using the bleach point tool of the Leica Software. Confocal and STED images of fixed samples were acquired with a Leica TCS SP8 gSTED 3X with 775nm STED laser (Leica Microsystems, Mannheim).

### SD length analysis

SD length was measured using a previously published macro.^10^ In a few cases, where the image contrast was insufficient to use the macro, the SD length was measured manually using FIJI.

### Calcium measurements

Calcium levels in podocytes were measured using FIJI and analyzed as previously published.^8^

### Statiscs

For statistical analysis of mean fluorescence intensities and SD length per area we used one-way ANOVA with Tukey’s multiple comparisons test. No data points were excluded. P values <0.05 were considered statistically significant.

## Results

We established a protocol for co-imaging of mouse glomeruli of Pod:cre-GCaMP3 mice, expressing the calcium indicator GCaMP3 exclusively in podocytes, with correlative multiphoton (MP) and STED microscopy. Prior to imaging, a triangle-shaped piece of tissue was cut from AKS at the upper left corner, to ensure identical slice orientation in both imaging modalities. During MP imaging, burn marks were positioned in proximity to glomeruli as reference points and large three-dimensional tile scans were acquired. These were used to measure the mean GCaMP3 fluorescence of glomeruli, as a readout for calcium levels in podocytes.^8^ To ensure tissue integrity and exclude tissue damage caused by cutting, AKS were incubated with propidium iodide and glomeruli with positive podocyte nuclei excluded from analysis. After this, AKS were fixed and processed for STED imaging as described in the method section. The previously imaged area was acquired again in three-dimensional image stacks, in which the added reference points allowed mapping of each glomerulus in the two data sets. Finally, STED images of the nephrin signal were acquired to visualize and quantify SD morphology (Fig. 1).

**Figure 1:**
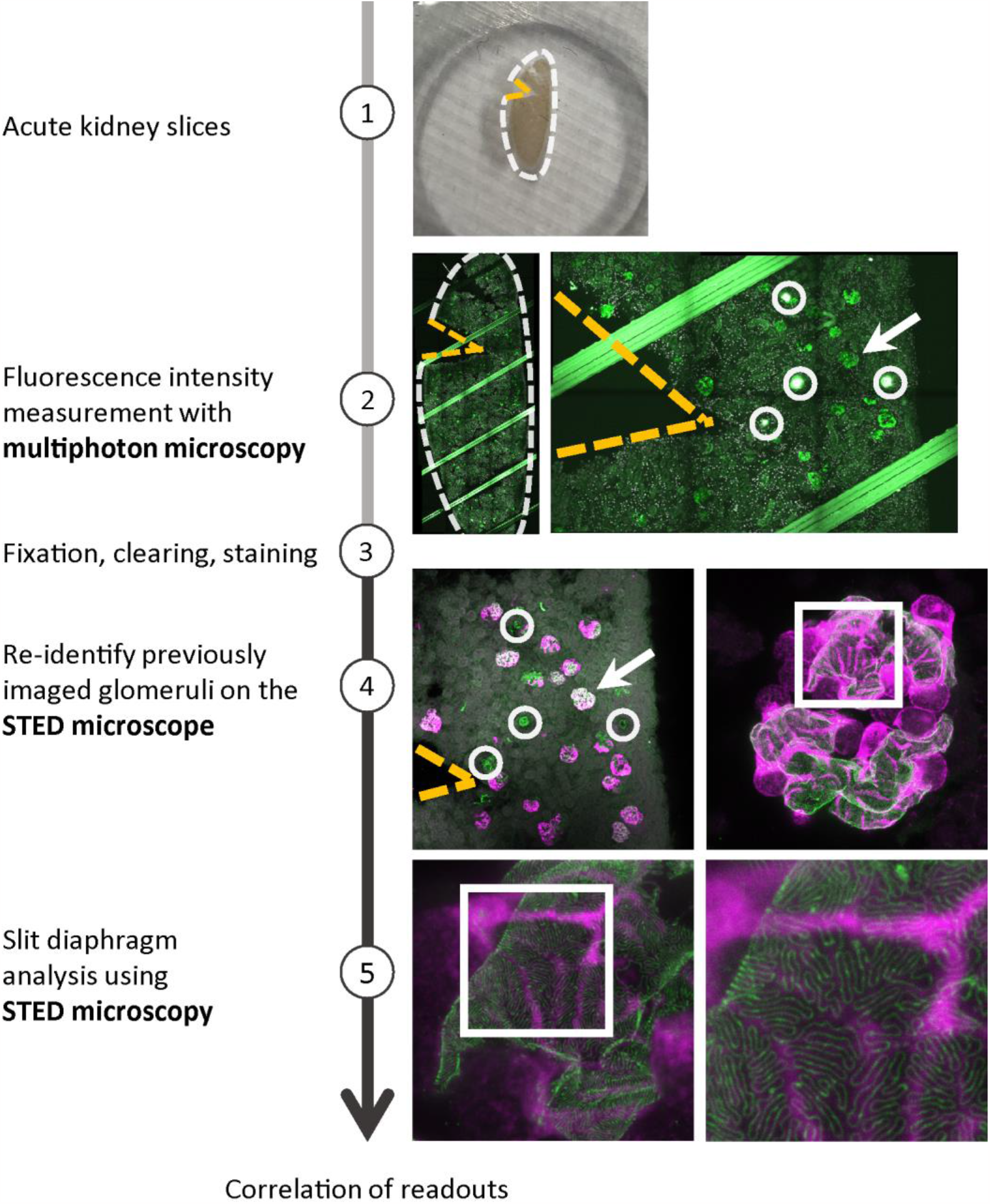
Workflow for co-imaging of murine glomeruli using multiphoton and STED microscopy. Overview of sample preparation and imaging steps. The microscope used is indicated on the left. 1 - AKS are marked by cutting a triangular shape at the upper left corner. 2 – Intensity measurements with multiphoton microscopy with burn marks as reference points (circles) and surrounding glomeruli (arrow). Green channel shows endogenous GCaMP3 fluorescence, propidium iodide staining in white. 3 – Fixation, clearing and staining of slices with goat anti-nephrin primary antibody, donkey anti-goat Alexa-594 secondary antibody and GFP-Alexa647. 4 – Confocal images with reference points (circles) and previsouly imaged glomeruli (arrows). 5 - STED images. Anti-GFP staining in magenta and anti-nephrin in green.

Comparison of the image stacks before and after processing confirmed the conserved morphology and validated the ability to study the same glomerulus with both imaging modalities (Fig. 2, Suppl. Video 1). We applied this protocol to image control animals (Fig. 2a) and mice injected with NTS in two doses: low (8 μl/g) and high dose (11 μl/g) (Fig. 2b). In all groups, we were able to re-identify the same podocytes within one glomerulus (Suppl. Video 1). Animals of both NTS groups showed significantly elevated calcium levels compared to control, but there was no significant difference between the two disease groups (Suppl. Fig. 1b). In both NTS groups, the SD length was significantly reduced compared to control. Between both disease groups the SD length was significantly lower in high dose than in low dose NTS animals, indicating damage to the podocyte architecture in a dose-dependent manner (Suppl. Fig. 1c). Of note, in both NTS groups, automatic and manual analysis of SD length in a subset of samples failed due to complete loss of a visible SD structure caused by extensive damage. The SD length was in these cases set to zero.

**Figure 2:**
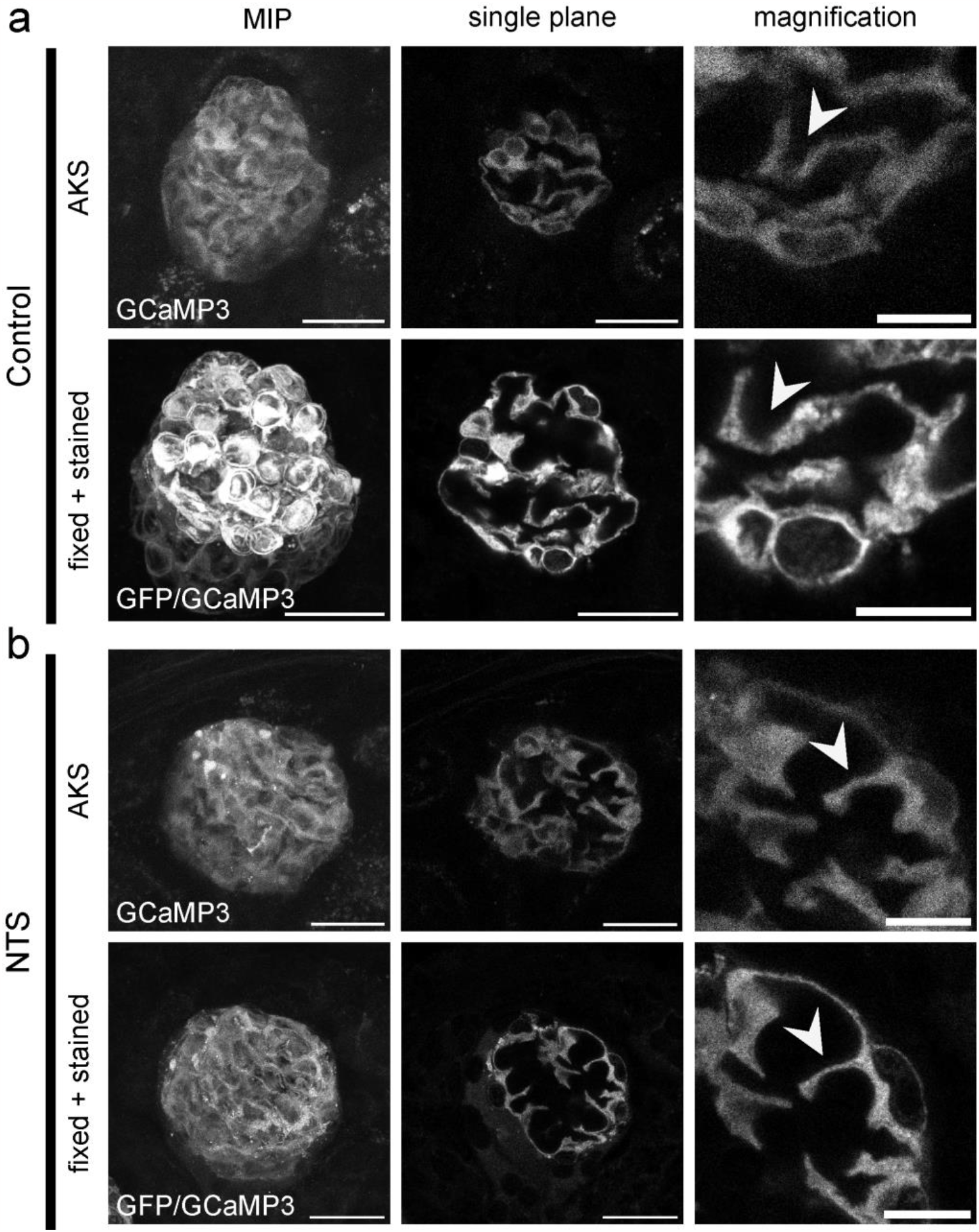
Reidentification of single mouse glomeruli and individual podocytes in AKS and after fixation and staining. Images of a control animal (a) and a NTS treated mouse (b). Images show same glomerulus – from left to right – maximum intensity projection (MIP) of whole glomerulus (left), single plane view (middle) and magnification of single podocytes (right). Endogenous GCaMP3 fluorescence for multiphoton images (AKS) and GFP/GCaMP3 signal for confocal images (fixed+stained). Arrowheads indicating single podocytes. Scale bar MIP & single plane – 25 μm; magnification – 10 μm.

Finally, we plotted intracellular calcium levels against the SD length for each glomerulus (Fig. 3a), which showed no correlation. However, the plot shows that 46/76 of glomeruli of low dose NTS mice had calcium levels exceeding the maximum levels of controls at the time point of measurement while all showed a reduction of SD length. In the high dose NTS group, only 19/36 of glomeruli had a measurable SD length. Representative images of SD morphology and GCaMP3 fluorescence for selected glomeruli from the different groups are shown in Figure 3b. In conclusion, we found that SD damage increases with higher NTS doses, while intracellular calcium levels revealed a large variance and showed no significant correlation with the extent of SD damage.

**Figure 3:**
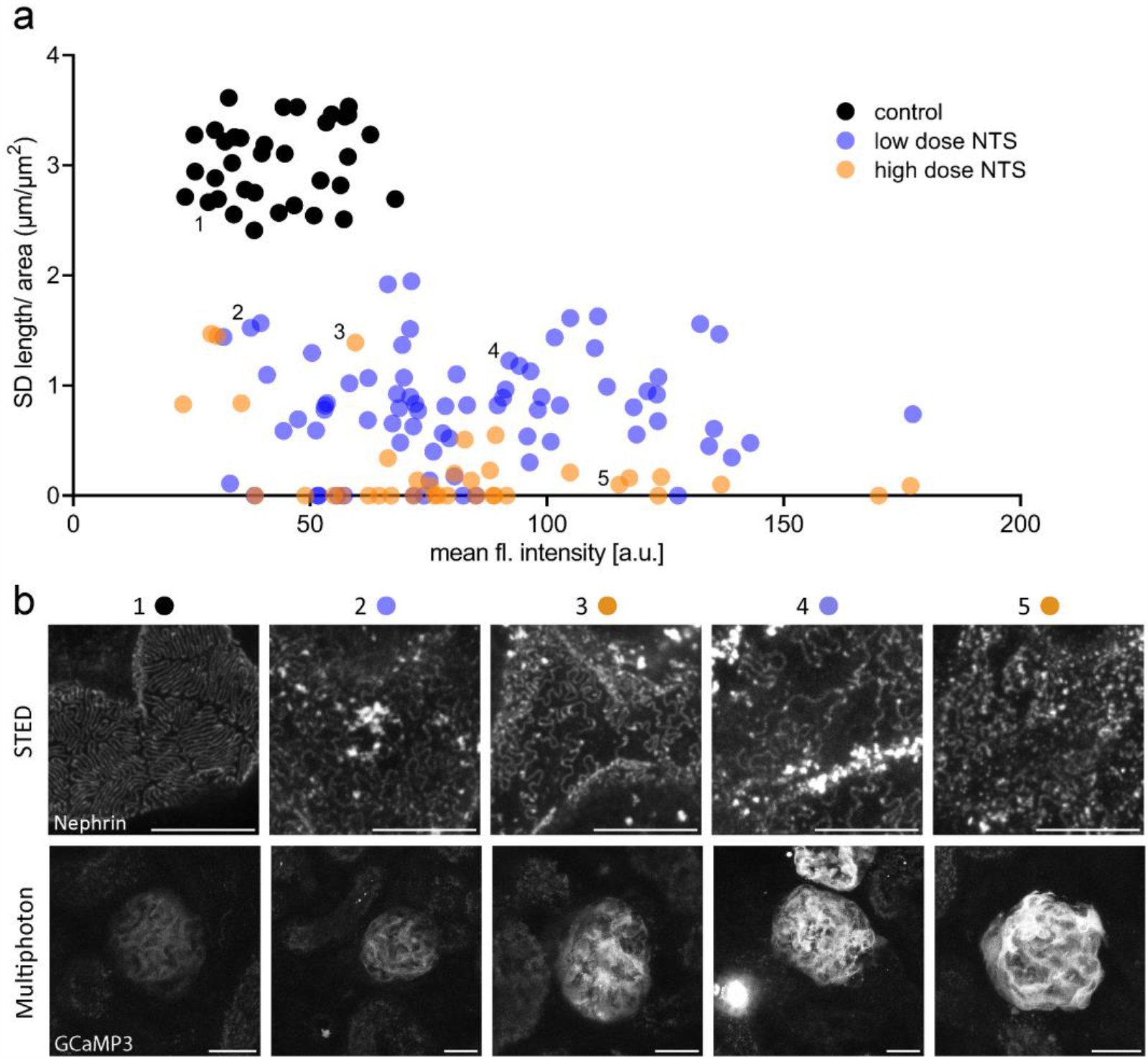
Correlation of intracellular calcium levels and podocyte slit diaphragm ultrastructure. (a) Correlation of mean fluorescence intensities of GCaMP3 and slit diaphragm length per area for individual glomeruli. (b) Representative MIP images of slit architecture acquired using STED imaging (upper panels) and GCaMP3 fluorescence in multiphoton microscopy (lower panels) of the same glomerulus. Symbols above panels show respective position in the quantification in (a). The following mice/samples were analyzed: Control: n=35 glomeruli of 3 mice; low dose NTS: n=76 glomeruli of 4 mice; high dose NTS: n=36 glomeruli of 3 mice. Scale bar STED – 5 μm; multiphoton images – 25 μm; MIP – max. intensity projection.

## Discussion

For the first time, our protocol allows visualization of functional information and nanoscale morphological information on the level of individual glomeruli. Using AKS and adding reference points to the tissue by laser damage enabled the identification of the same glomerulus in the second imaging modality.

The constantly increasing availability of genetically encoded fluorescent sensor proteins for *in vivo* imaging and immunofluorescence labeling of proteins of interest make our co-imaging workflow suitable for a broad range of research questions. In this study, we visualized calcium levels in podocytes in health and disease and correlated those with the integrity of the SD. Adding a perfusion pump to exchange buffers would allow for triggering and recording of calcium responses to a substance of choice in a high temporal resolution. This would enable the correlation of calcium signaling to subsequent changes in podocyte ultrastructure by STED microscopy.

Our data shows, that all glomeruli of control animals exhibited low intracellular calcium levels as previously shown *in vivo*.^8^ After induction of NTS nephritis, both groups showed an increase of the mean fluorescence, but with a large variation between glomeruli probably due to the heterogenous nature of the NTS model. In half of glomeruli of animals receiving high dose NTS, nephrin abundance was diminished to such a degree that analysis of SD length by STED imaging was not possible (Fig. 3a). Without preselection of glomeruli by multiphoton imaging, glomeruli with complete disruption of the SD architecture would likely not be selected for analysis, as these glomeruli are absent of a nephrin SD pattern. Therefore, the published SD analysis protocol might underestimate the severity of alterations in podocyte architecture in very advanced cases of glomerular disease.^10^ Low dose NTS had a less pronounced effect on SD architecture as measured by a reduction in SD length (Fig. 3 or Suppl. Fig. 1) compared to high dose NTS. Even though our method clearly showed a dose-dependent effect of NTS on the integrity of the SD, we could not observe a correlation between calcium and a reduction in SD length. While intracellular calcium levels can change rapidly, morphological changes to the SD take far longer to form and resolve. Therefore, calcium measurement at a single time point might not be sufficient to draw conclusions about the extent of glomerular damage, even if mean calcium levels increase in disease.

In conclusion, we present a protocol for co-imaging AKS with multiphoton and STED microscopy in order to correlate functional and structural information on the level of individual glomeruli and possibly even single glomerular cells.

## Acknowledgements

Calcium multiphoton and STED imaging experiments were designed by JB-L, AH and MJH. Experiments were performed by EW, NR and JB-L. Data analysis was carried out by EW and DU-J. The manuscript was written by EW, JB-L and MJH. TB and MJH provided guidance and resources. The authors acknowledge the help of the Cologne Excellence Cluster on Cellular Stress Response in Aging-Associated Diseases Imaging Facility (Dr. Christian Jüngst and Dr. Astrid Schauss).

## Data sharing statement

The imaging data acquired by STED and multiphoton microscopy analyzed in this study is available in the Figshare repository at https://figshare.com/s/79be57e74e3fe693876b with the reserved DOI: 10.6084/m9.figshare.24118053.

