## Supplemental Figure 1 for "Correlation of calcium multiphoton with ultrastructural STED imaging of the slit diaphragm in the same glomerulus"

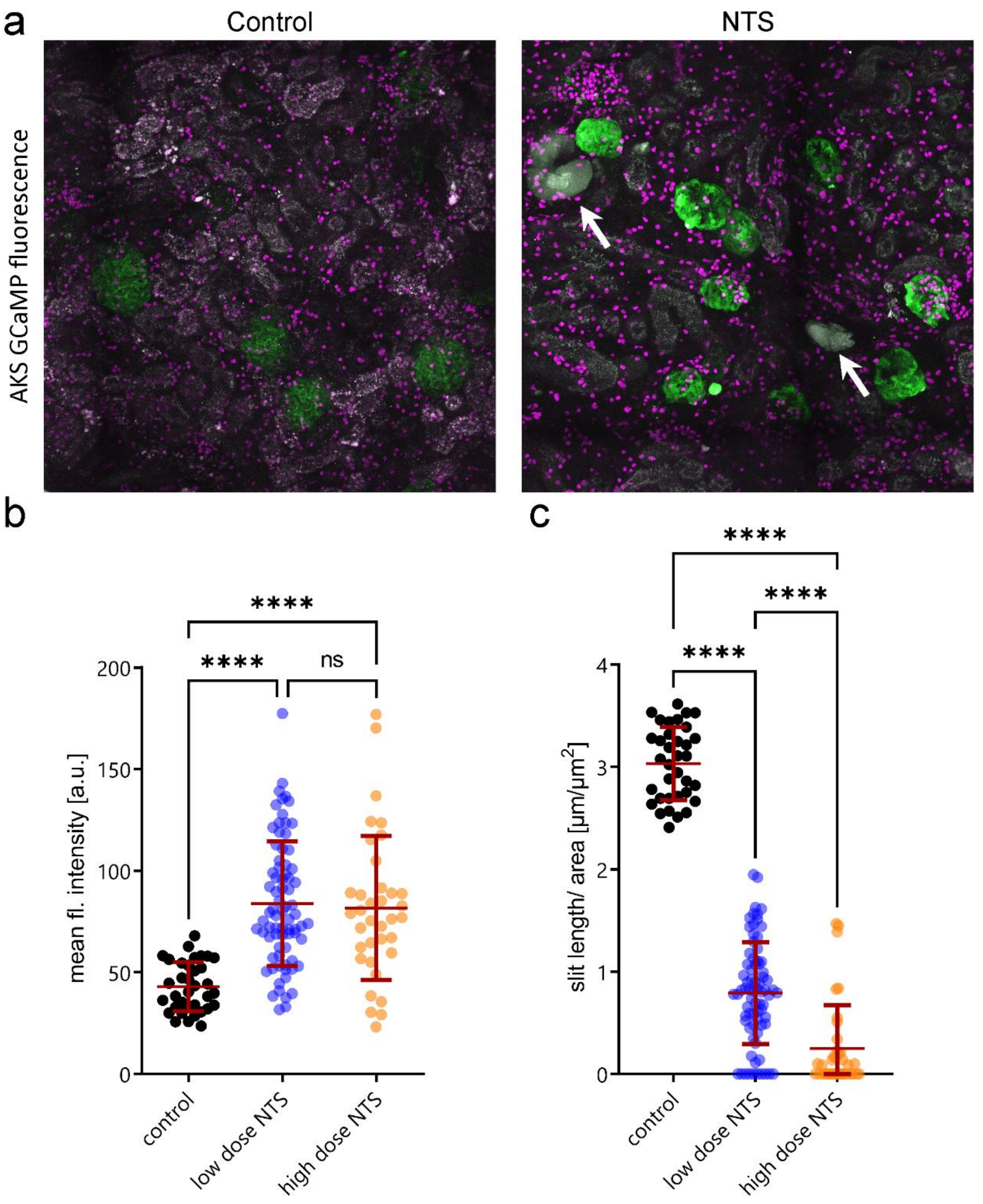


**Supplemental Figure 1: GCaMP3 fluorescence as an indicator for podocyte $[Ca^{2+}]_i$ in acute murine kidney slices.** (a) Representative MIP images of podocyte specific GCaMP3 fluorescence in AKS of control and NTS (high dose) treated mice. GCaMP3 expression in podocytes (green), propidium iodide staining labeling dead cells (magenta). Protein casts in tubules in diseased animals indicated by arrows. Acquired using multiphoton microscopy. (b) Quantification of mean fluorescence intensities of glomeruli for control, low dose NTS (8 µl/g) and high dose NTS (11 µl/g BW) mice. Control mean: 42.9 ±12.0; low dose NTS mean: 83.8 ±30.4; high dose NTS mean: 81.6 ±34.9. (c) Quantification of SD length per area of single foot processes of healthy control, low dose NTS (8 µl/g,) and high dose NTS (11 µl/g) treated animals. Glomeruli in which no SD could be measured are set to value = 0. Control mean: 3.0±0.4; low dose NTS mean: 0.8 ±0.5; high dose NTS mean: 0.3 ±0.4. Images were acquired using STED microscopy and analyzed with a FIJI macro or manually. The following mice/samples were analyzed: Control: n=35 glomeruli of 3 mice; low dose NTS: n=76 glomeruli of 4 mice; high dose NTS: n=36 glomeruli of 3 mice. Statistical analysis: ordinary one-way ANOVA with Tukey's multiple comparisons test; p-value ****<0.0001. Red line represents mean + standard deviation. AKS – acute kidney slices; MIP – max. intensity projection; NTS – Nephrotoxic serum; SD – slit diaphragm; ns – not significant.

**Supplemental Video 1: Structural overview of the same glomerulus imaged with a multiphoton microscope (left) and confocal settings on a STED microscope after processing (right).** The left stack shows a glomerulus of a healthy mouse expressing GCaMP3 in podocytes acquired by multiphoton imaging. The right stack shows the same glomerulus after processing (fixation and staining) acquired with confocal imaging settings using a STED microscope. Comparison of both image stacks clearly confirms positive identification of the same glomerulus in both imaging modalities.
